## Supplementary figures for "Circadian clock mechanism driving mammalian photoperiodism"

Wood et al

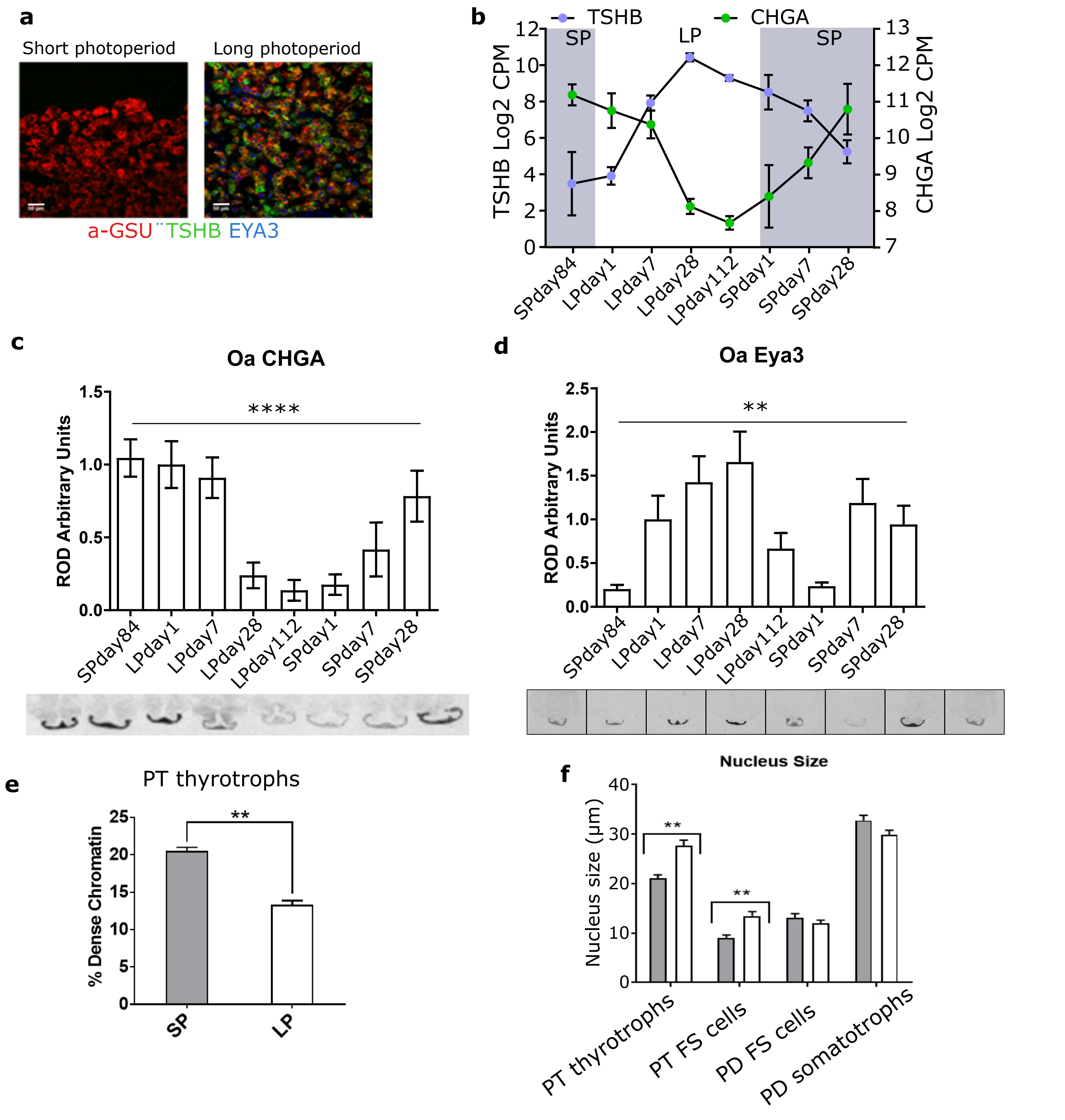


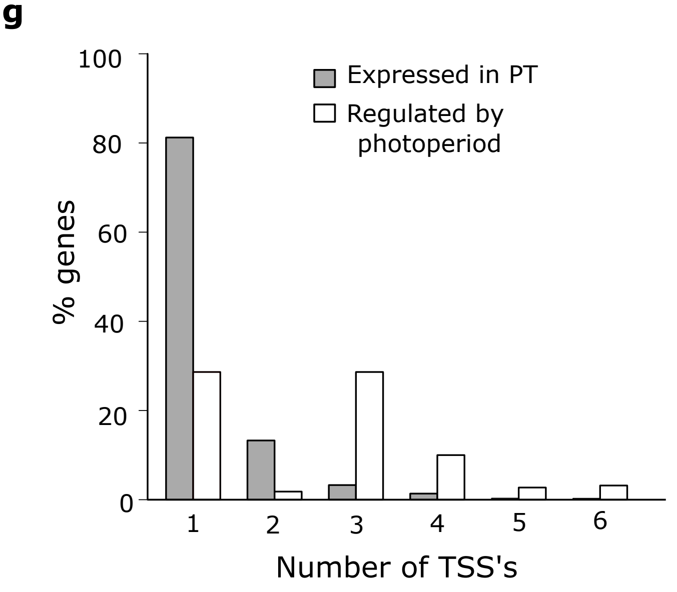


Supplementary Figure 1

**Supplementary Fig. 1: Photoperiod dependent epigenetic regulation of transcription in the Pars tuberalis**

a. Triple immunofluorescence showing expression of aGSU (red), TSHb (green), and EYA3 (blue) in the PT on SP day 28 and LP day 28. Scale bars, 50 um.

b. RNAseq log2 counts per million (CPM) of TSHb (purple) and CHGA (green) over the experiment. Grey shading represents SP sampling points. Error bars represent the SEM.

c. In situ hybridization and quantification for CHGA mRNA at SP day 84, LP day 1, 7, 28, 112, SP day 1, 7, 28. Representative images are shown (n = 3). Error bars represent the SD. Statistical analysis by one-way ANOVA confirms there is a significant seasonal change. **** = p-value less than 0.00005.

d. As in c for EYA3 mRNA . ** = p-value less than 0.005.

e. PT thyrotroph chromatin density at LP and SP day 28, given as the percentage of dense chromatin contained within the nucleus relative to nucleus size. N=3 individuals from SP and LP. 40 nuclei were measured per group. T-test was used to assess statistical significance. **= FDR less than 0.00001

f. PT thyrotroph, PT follicular stellate (FS) cell, pars distalis (PD) FS cells and PD somatotroph nucleus measurements (um) at LP and SP day 28. N=3 individuals from SP and LP. 40 nuclei were measured per group. T-test was used to assess statistical significance. **= FDR less than 0.00001

g. Percentage of genes with a given number of transcription start sites in the genomic background (all >0 log2CPM expressed genes) of the pars tuberalis (grey bars) as compared to all seasonally differentially expressed genes (white bars).

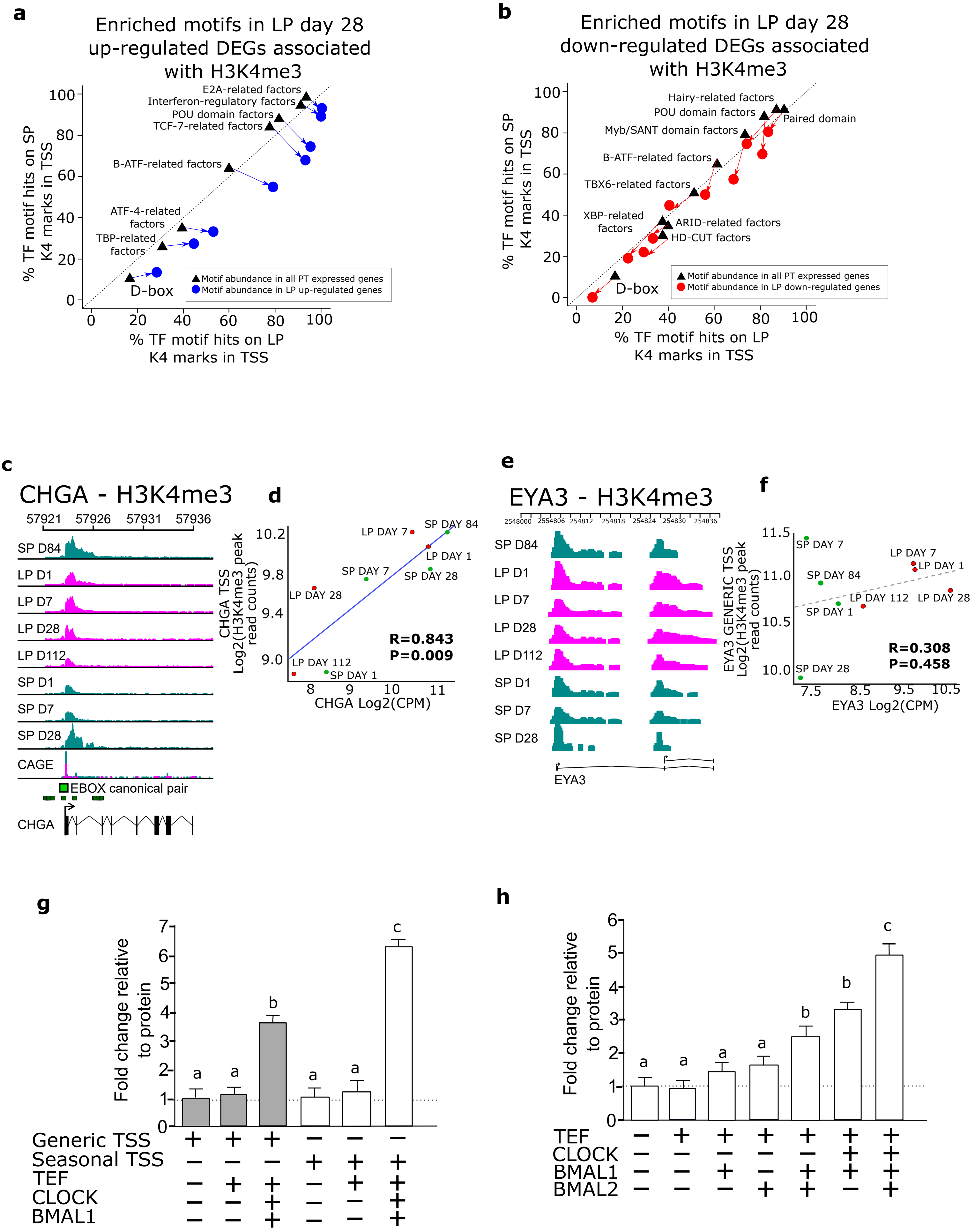


Supplementary Figure 2

**Supplementary Fig. 2: Epigenetic regulation of a seasonal E-box enriched promoter in EYA3**

a. Plot of all over-represented motifs in the active promoters of LP day 84 upregulated genes. The axis plot the percentage of inferred active promoters, containing one or more observed motifs, for a given cohort of genes. Active promoters are defined as contiguous H3K4me3 marked regions within 100bp of a TSS. The black triangles represents the motif abundance on active promoters in all the genes expressed (>0 CPM) in the pars tuberalis (PT) and the percentage of their active promoters containing a binding motif for LP (x-axis) and SP (y-axis) H3K4me3 environments. The circles are the motif coverage of activate promoters in differentially up-regulated LP genes (FDR < 0.05; fishers two-way exact test).

b. As in G for the over-represented motifs in the TSS of LP down-regulated genes when compared to SP day 84.

c. ChIP-seq tracks for CHGA gene H3K4me3 peaks for the whole experiment. Chromsome 2 region is shown. Pink represents samples in long photoperiod and marine green represents short photoperiod. Solid green arrows are canonical Ebox motifs. Blue arrows are D-box motifs. Gene schematic is also shown.

d. Correlation plot for CHGA downstream TSS log2 H3K4me3 peaks from ChIPseq versus CHGA log2 counts per million (CPM) from RNA-seq. Red symbols are LP sampling points, green are SP sampling points. Correlation coefficient R is shown. R=0.843, p-value=0.009.

e. As in A for EYA3 gene. Chromsome 2 region is shown.

f. As in B for EYA3 up-stream TSS. R=0.368, p-value=0.458.

g Transactivation of the EYA3-upstream-TSS-luc reporter versus the EYA3-downstream-TSS-luc reporter by TEF, CLOCK and BMAL1. The experiment was repeated 4 times (n=4 per experiment), plot displayed is a representative result. A one-way ANOVA was performed on each individual experiment using Tukey's multiple comparisons test. Different letters indicate significant differences between groups (P < 0.01). Error bars SEM.

h Transactivation of the EYA3-downstream-TSS-luc reporter by TEF, CLOCK, BMAL1 and BMAL2. The experiment was repeated 4 times (n=4 per experiment), plot displayed is a representative result. A one-way ANOVA was performed on each individual experiment using Tukey's multiple comparisons test. Different letters indicate significant differences between groups ($P < 0.01$). Error bars SEM.

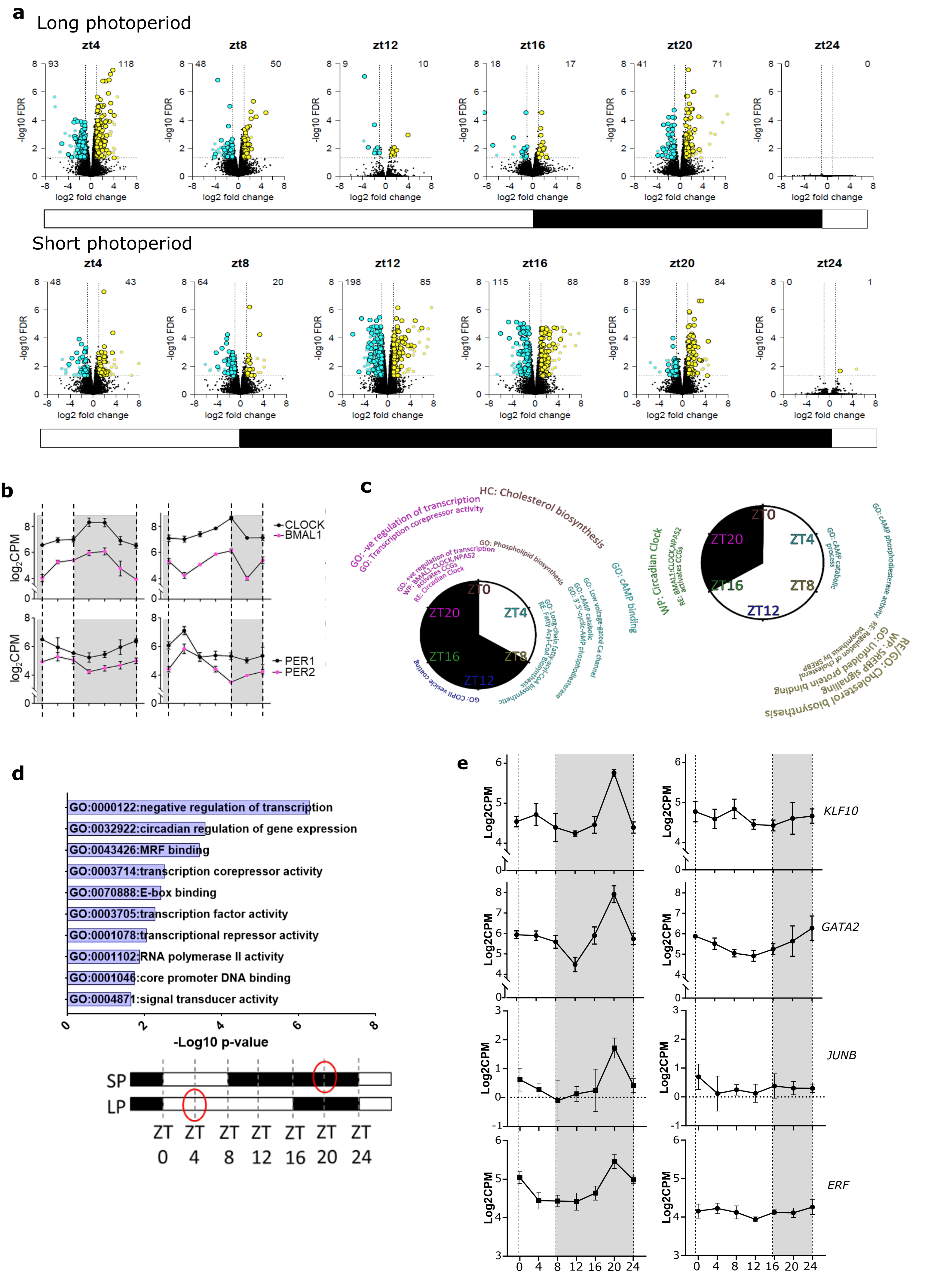


Suppelmentary Figure 3

**Supplementary Fig. 3: Diurnal gene expression analysis show enrichment for repressors at SP ZT20**

a. Volcano plots showing the number of genes up (yellow) and downregulated (blue) across the day in SP and LP. All comparisons were pairwise against ZT0 pairwise comparisons. ZT24 is included to show the consistency between timepoints. ZT timepoints are indicated above each plot and the light dark transistion is shown below. Numbers on the plots are the number of significantly up or down regulated genes in that pairwise comparison (Supplementary Data 6).

b. RNA-seq log2 CPM plots. Light dark bars shown a the bottom and indicated on the graph by dotted line and grey shading. Error bars are SD. Statistical significances are in (Supplementary Data 6).

c. Enrichment term analysis using CPDB and radial plots of ZT time against -log10 pvalue (terms further out have lower pvalues) to indicate the functional enrichments at each peak phase.

d. TopGO analysis of genes differentially expressed between SP ZT20 and LP ZT4, i.e. 12 hours after lights off. Enrichment is shown as –Log10 p-value.

e. RNA-seq log2 CPM plots. Light dark bars shown a the bottom and indicated on the graph by dotted line and grey shading. Error bars are SD. Statistical significances are in (Supplementary Data 6).

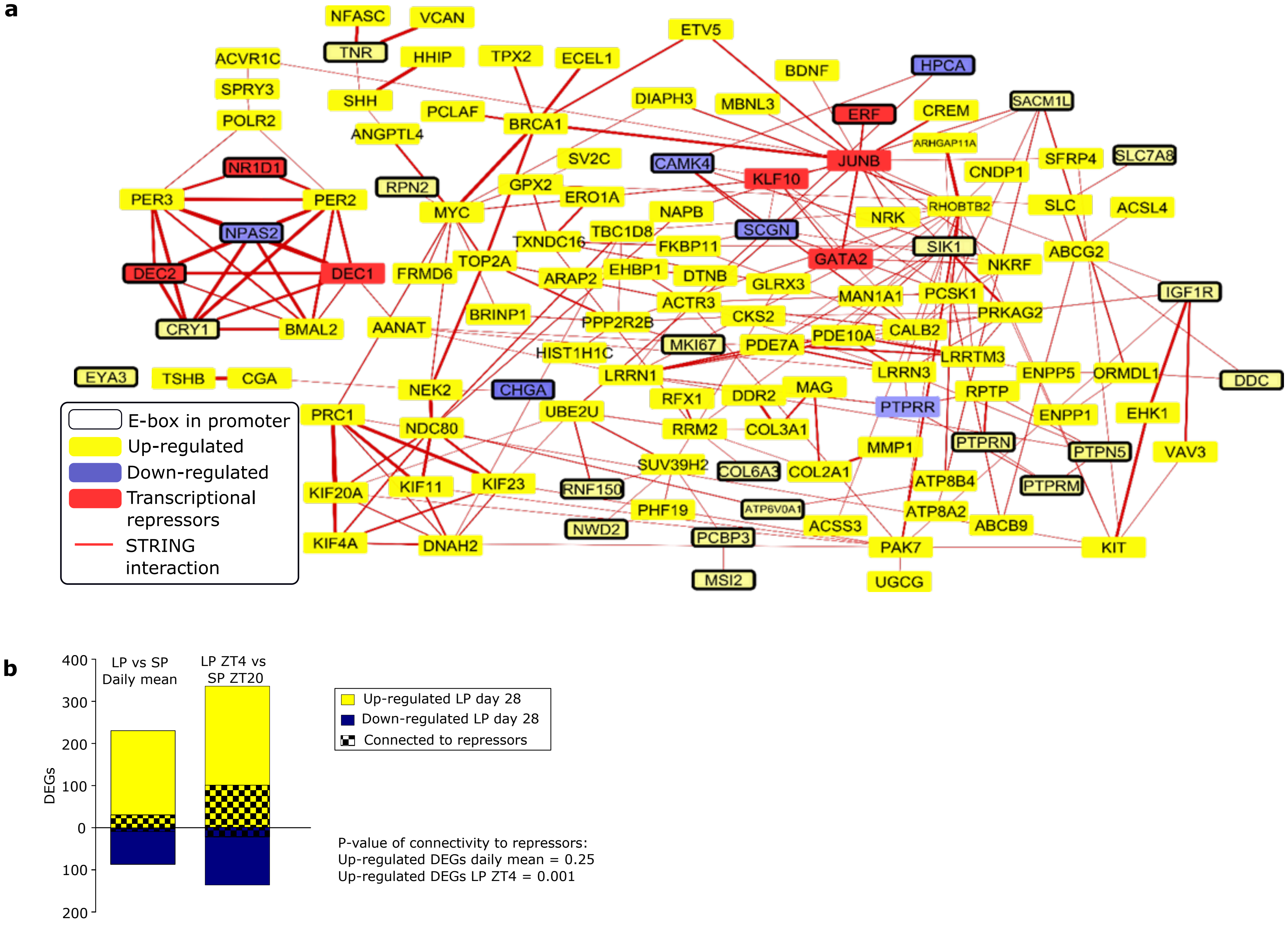


Supplementary Figure 4

**Supplementary Fig. 4: STRING analysis reveals that the SP ZT20 repressors are high connected to the LP ZT4 transcriptome**

a. Repressor genes up-regulated at ZT20 in SP (red boxes) were analysed for their connectivity, and therefore interaction with the genes up-regulated at ZT4 in LP (Yellow boxes and down-regulated (blue boxes). A high degree of proteinprotein intereaction (PPI) connectivity was found to the up-regulated genes (see red lines on network). Black boarders on gene boxes indicate presence of Eboxes in the promoter.

b. Number of differentially expressed genes (DEGs), up-regulated (yellow) and down-regulated (blue) for a daily mean (all 24 hour timepoints) between SP vs LP on day 28, compared to LP ZT4 vs SP ZT20 contrast. Connectivity via protein-protein intereactions (PPI), defined by STRING to transcriptional repressors is indicated by the checkered shading (also represented in Fig. 4a). The significance of enrichment (fishers two-way exact test) for PPI connectivity within the up-regulated vs down-regulated genes is shown.

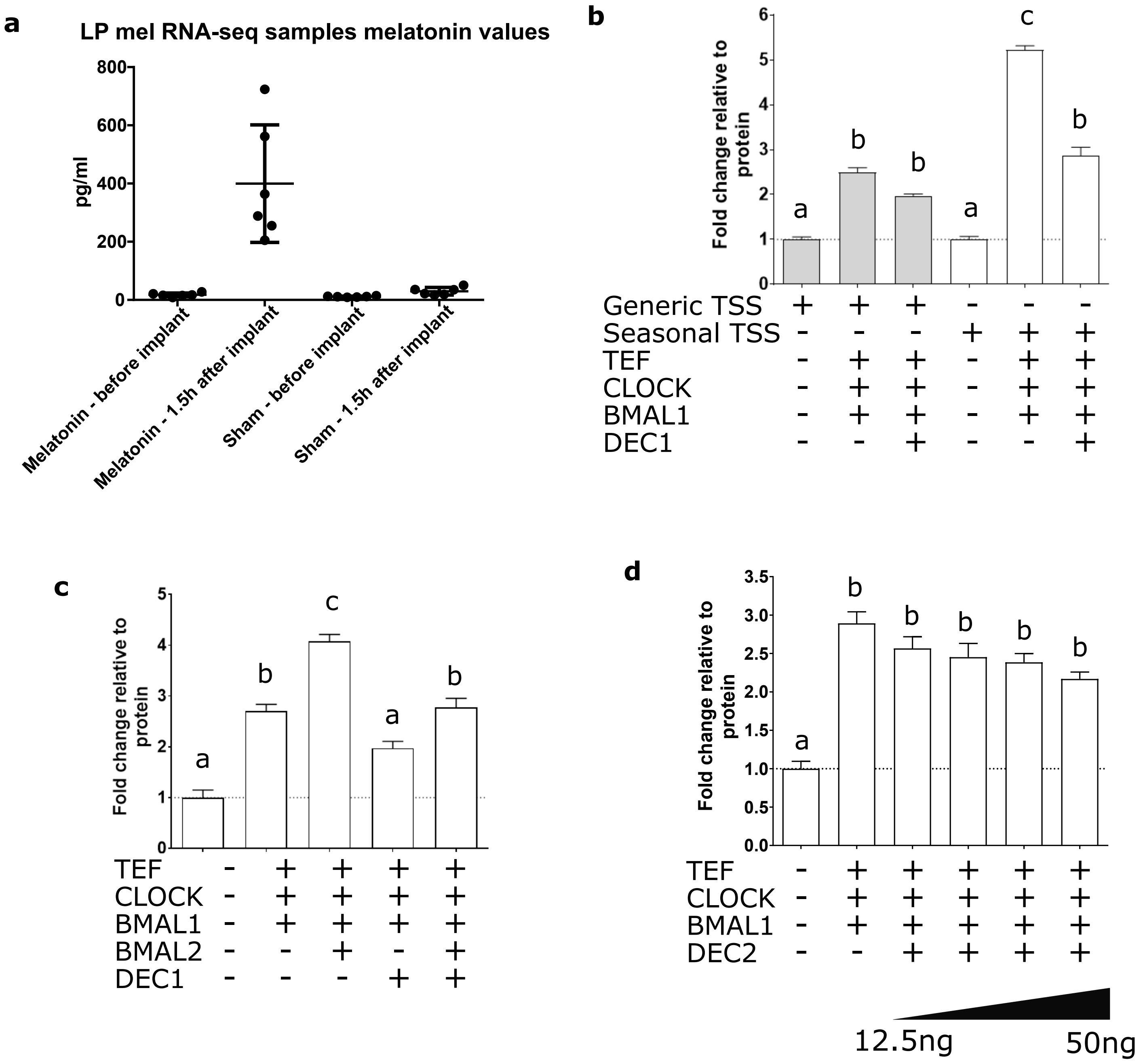


Supplementary Figure 5

**Supplementary Fig. 5: Melatonin implant validation and DEC1 repression**

a. Melatonin concentration after implantation compared to sham and pre – implantation. Values are pg/ml. Mean and SD shown.

b. Transactivation of the EYA3-downstream-TSS-luc reporter versus EYA3-upstream-TSS-luc reporter by TEF, CLOCK, BMAL1 and BMAL2, and the effect of DEC1. The up-stream promoter is not significantly repressed by DEC1 but the downstream TSS activation is significantly suppressed. The experiment was repeated 4 times (n=4 per experiment), plot displayed is a representative result. A one-way ANOVA was performed on each individual experiment using Tukey's multiple comparisons test. Different letters indicate significant differences between groups ($P < 0.01$). Error bars SD.

c. Transactivation of the EYA3-downstream-TSS-luc reporter by TEF, CLOCK, BMAL1 and BMAL2, and the suppression by DEC1. The experiment was repeated 4 times (n=4 per experiment), plot displayed is a representative result. A one-way ANOVA was performed on each individual experiment using Tukey's multiple comparisons test. Different letters indicate significant differences between groups ($P < 0.01$). Error bars SEM. d. Transactivation of the EYA3-downstream-TSS-luc reporter by TEF, CLOCK and BMAL1, and the effect of DEC2 (12.5 to 50ng). The experiment was repeated 4 times (n=4 per experiment), plot displayed is a representative result. A one-way ANOVA was performed on each individual experiment using Tukey's multiple comparisons test. Different letters indicate significant differences between groups ($P < 0.01$). Error bars SD.

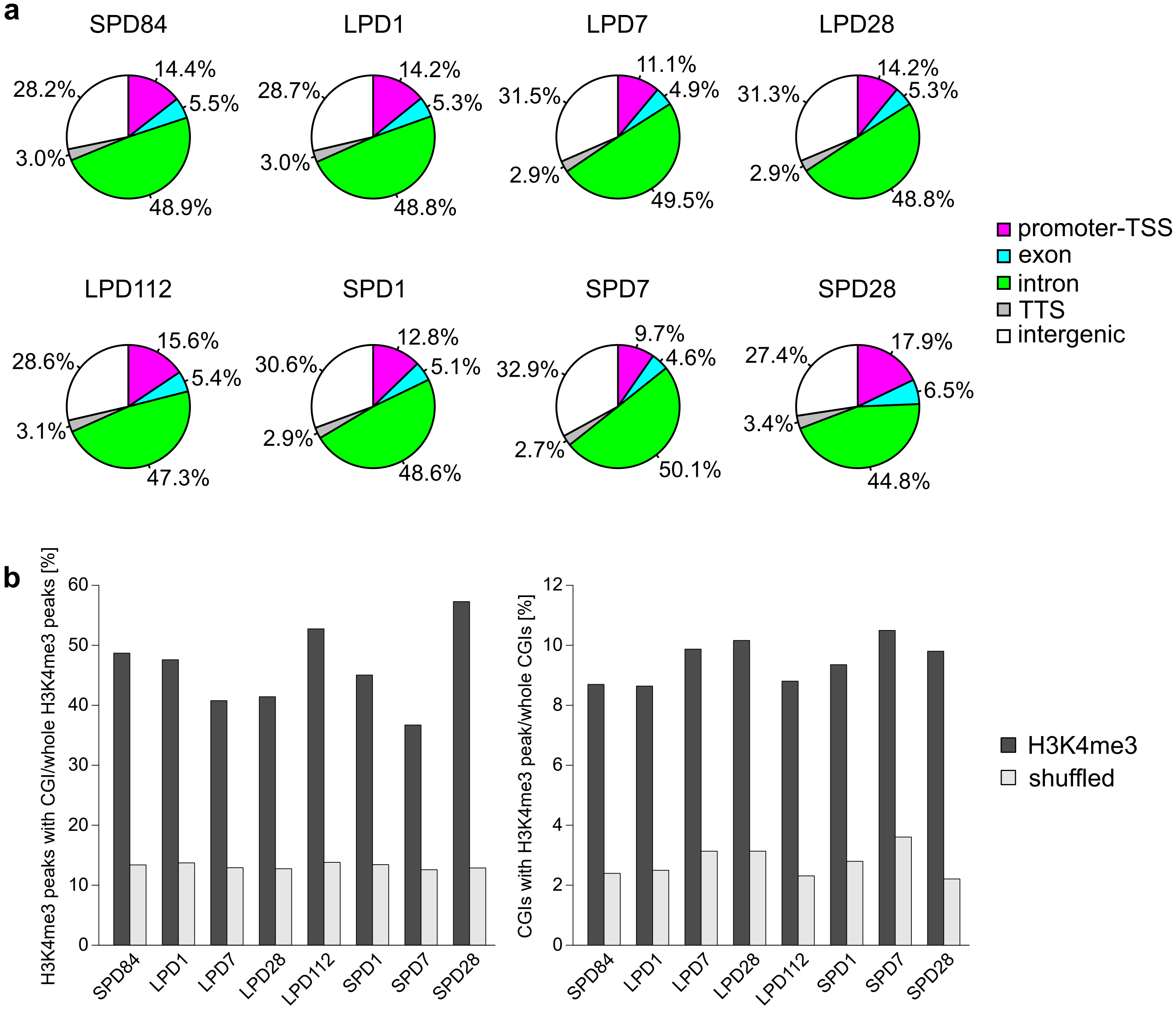


Supplementary Figure 6

**Supplementary Fig. 6: Quality check of H3K4me3 ChIP-seq.**

a. Pie charts revealing distributions of H3K4me3 peaks on each genomic feature. Peaks of promoter-TSS were located on ±1000 bp from TSS.

b. Bar plots revealing percentages of H3K4me3 peaks co-localised with CGIs (left) and CGIs co-localised with H3K4me3 peaks (right) on the sheep genome. Black is observed H3K4me3 peaks in PTs and grey is randomly shuffled peaks with the same fragment sizes as negative controls.

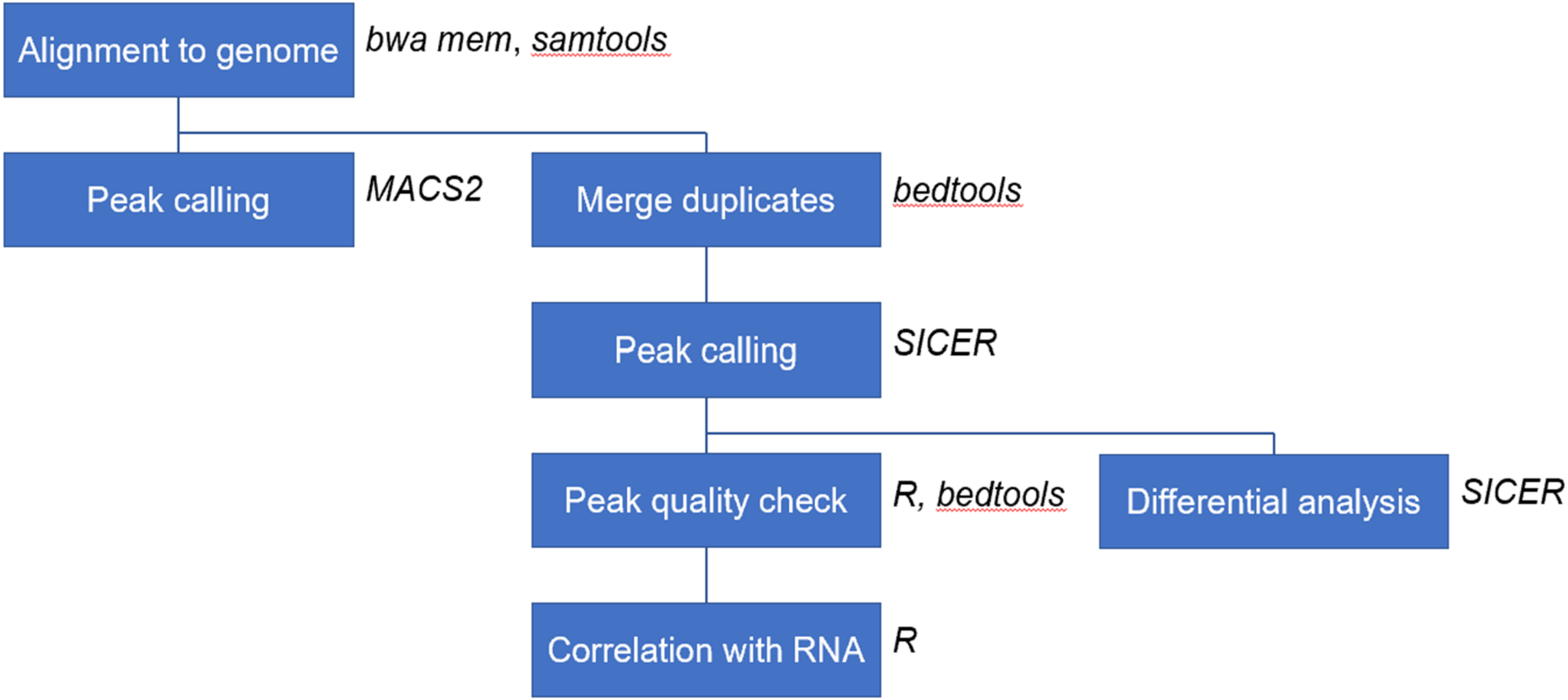


Supplementary Figure 7

**Supplementary Fig. 7:** Overview the analysis workflow for ChIP-seq
