## Supplementary data (Tables) descriptions for "Circadian clock mechanism driving mammalian photoperiodism"

Wood et al

List of Supplementary Data

Supplementary Data 1: Seasonal epigenetic changes, MACs peak calling ChIP-seq H3k4me3

Supplementary Data 2: Seasonal epigenetic changes, SICER differential peak analysis for H3K4me3

Supplementary Data 3: Seasonal gene expression, pairwise contrasts for RNA-seq

Supplementary Data 4: CAGE transcription start site clusters and their relative seasonal abundance

Supplementary Data 5: RNA-seq expression for known H3K4me3 modulators

Supplementary Data 6: Diurnal gene expression, statistical analysis of 24 hour profiles

from RNA-seq
